# A classification framework for *Bacillus anthracis* defined by global genomic structure

**DOI:** 10.1101/675249

**Authors:** Spencer A. Bruce, Nicholas J. Schiraldi, Pauline L. Kamath, W. Ryan Easterday, Wendy C. Turner

## Abstract

*Bacillus anthracis*, the causative agent of anthrax, is a considerable global health threat affecting wildlife, livestock, and the general public. In this study whole-genome sequence analysis of over 350 *B. anthracis* isolates was used to establish a new high-resolution global genotyping framework that is both biogeographically informative, and compatible with multiple genomic assays. The data presented in this study shed new light on the diverse global dissemination of this species and indicate that many lineages may be uniquely suited to the geographic regions in which they are found. In addition, we demonstrate that plasmid genomic structure for this species is largely consistent with chromosomal population structure, suggesting vertical inheritance in this bacterium has contributed to its evolutionary persistence. This classification methodology is the first based on population genomic structure for this species and has potential use for local and broader institutions seeking to understand both disease outbreak origins and recent introductions. In addition, we provide access to a newly developed genotyping script as well as the full whole genome sequence analyses output for this study, allowing future studies to rapidly employ and append their data in the context of this global collection. This framework may act as a powerful tool for public health agencies, wildlife disease laboratories, and researchers seeking to utilize and expand this classification scheme for further investigations into *B. anthracis* evolution.

## INTRODUCTION

*Bacillus anthracis* is a Gram-positive, rod-shaped bacterium and world-wide clonal zoonotic pathogen, best known as the etiologic agent of anthrax. Bacterial infections of *B. anthracis* are primarily relegated to wild and domestic herbivores, with occasional spillover to humans (Carlson et al., 2018; de Vos et al., 2018). *B. anthracis* has also garnered global attention as a bioterrorism weapon due to the highly resilient qualities of its endospores, and their ability to be aerosolized for malicious purposes (Mugavero et al., 2015; Koehler & Popella, 2018). Virulence in *B. anthracis* is dependent on the chromosome, (> 5,200,000 base pairs (bp) in length), and two large plasmids: pXO1 (> 180,000 bp), and pXO2 (> 90,000 bp). The plasmid pXO1 harbors genes responsible for the anthrax toxin, while the pXO2 plasmid encodes for both capsule synthesis and degradation (Kolstø et al., 2009; Fasanella, 2013). Both plasmids are vital for *B. anthracis* pathogenicity, and the absence of either greatly reduces the potential virulence of the bacterium.

Researchers, public health institutions and livestock regulation agencies have largely relied on PCR-based methods such as multilocus variable number of tandem repeat analysis (MLVA) and chromosomal single nucleotide polymorphism (SNP) analysis for strain typing and differentiation (Keim et al., 2000; Ellerbrok et al., 2002; Van Ert et al., 2007a; Beyer et al., 2012; Ogawa et al., 2015).While these methods have the potential to provide insight into global phylogenetic relationships, they are limited in that they rely heavily on the finite number of samples used for initial marker selection, and may produce little information on plasmid genetic structure and/or differentiation (Eyre et al., 2013; Ranjbar et al., 2014).

In 2004, SNP discovery based on 5 genomes was used to identify approximately 3,500 chromosomal SNPs among a unique set of *B. anthracis* strains. Nearly 1,000 of these SNPs were then mapped across 27 diverse isolates, chosen based on MLVA analysis to develop the then current phylogenetic model for this species describing three major lineages: A, B, and C (Pearson et al., 2004). This work greatly increased our understanding of global diversity and evolution for *B. anthracis* in terms of phylogenetic structure. Later efforts that built on these data were used to develop a genotyping framework based on 12 chromosomal SNPs, enabling the classification of *B. anthracis* into the three previously recognized major lineages (A, B, and C), with further subdivision into 12 sub-lineages (Van Ert et al., 2007b). When examining the population genetic structure of this species in tandem with the sub-lineage classifications, results demonstrated classifications inconsistent with population genetic structure based on VNTR analysis in some instances, as well as a limited ability to discriminate isolates into lineages at the population level (Van Ert et al., 2007a). This framework has been incorporated into the canSNPer package for cladistic categorization and remains the current standard for *B. anthracis* classification across the globe (Lärkeryd et al., 2014; Van Ert et al., 2007a). These methods have been used to document genotypic diversity in anthrax outbreaks occurring from local to regional scales (Rume et al., 2016a; Rume et al., 2016b; Lienemann et al., 2018; Zhang et al., 2018).

More recently, whole genome sequencing (WGS) technology has helped to resolve local differences in *B. anthracis* strains extending the aforementioned genotyping scheme (Girault et al., 2014a; Girault et al., 2014b; Derzelle et al., 2015), but currently no system based on WGS data, using high resolution SNP detection, has been incorporated into a large-scale global framework, such as those produced for other monophyletic clades such as *Salmonella enterica, Mycobacterium tuberculosis*, and *Yersinia pestis* (Morelli et al., 2010; Coll et al., 2014; Wong et al., 2016), and no genotyping system defined by population genomic structure for this species currently exists.

In this study we developed a global genotyping system for *B. anthracis* using the WGS data of 356 *B. anthracis* isolates collected from 39 countries over the course of a century, available from the National Center for Biotechnology Information (NCBI) database (Table 1). We identified >14,500 chromosomal SNPs in the core genome, >580 SNPs in the pXO1 plasmid, and >320 SNPs in the pXO2 plasmid, all of which were used to construct comprehensive phylogenetic trees for each bacterial component. We then used this high resolution WGS dataset to re-define the global systematics for *B. anthracis* using current methods for population genomic analysis, establishing a robust framework for primary cluster and clade designation based on hierarchical genomic structure. These results not only reveal the global biogeographic structure of *B. anthracis* lineages, but may also act as a powerful tool for livestock and wildlife health surveillance, epidemiological studies, anti-terrorism investigations, and laboratory experiments involving *B. anthracis* and ultimately anthrax.

**Table 1.**
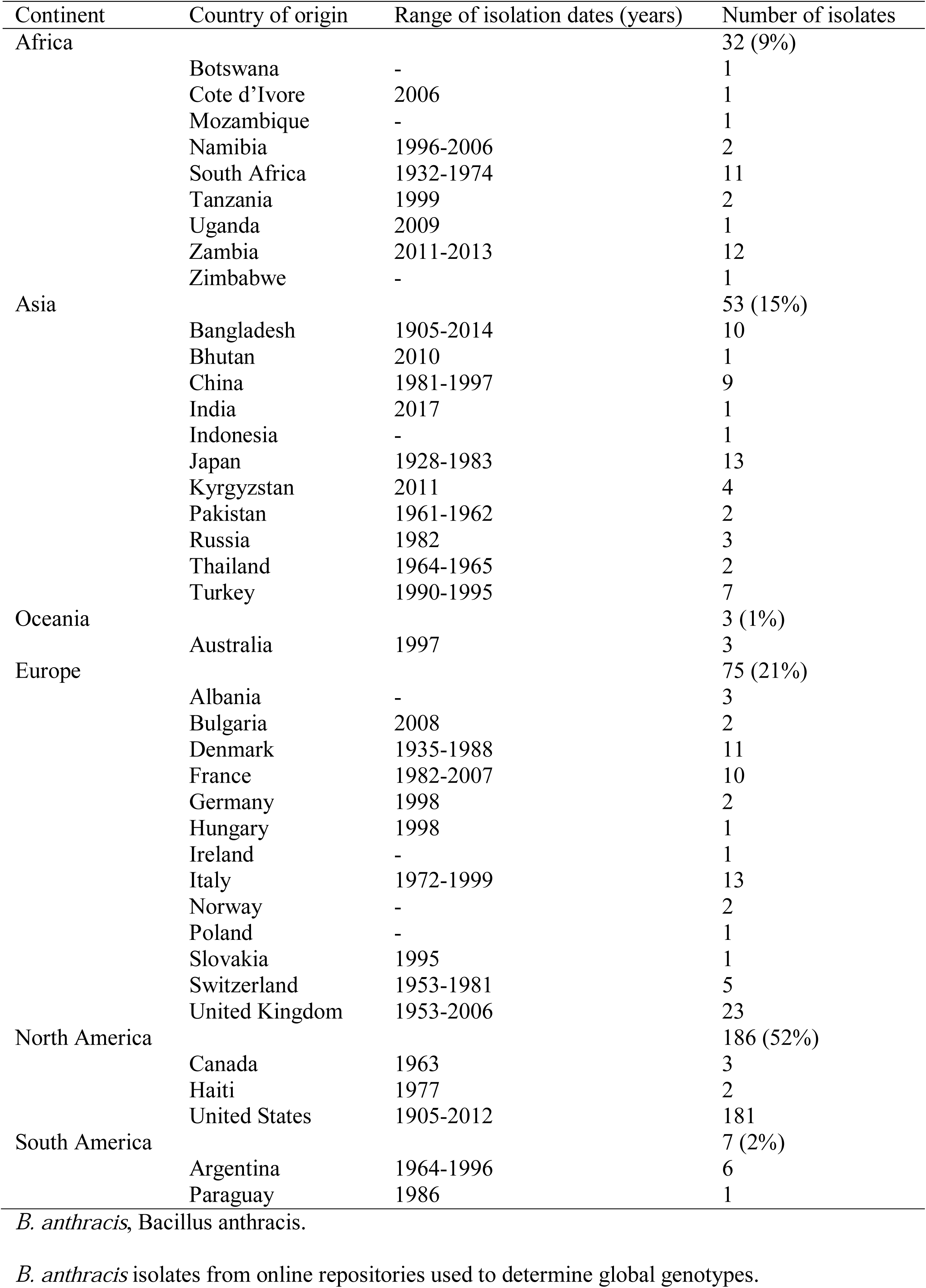
Summary of 356 *B. anthracis*.

## METHODS

### Bacterial isolates and WGS data

A total of 422 *B. anthracis* isolates collected from both natural hosts (wildlife and livestock) and directly from the environment were compiled for this study. Paired-end Illumina raw read data included in the analysis was freely available for download from public databases, and followed variable WGS protocols (S1 Dataset). Sample isolation dates range from 1905 through 2018, encompass 39 countries, and span 6 continents (Africa, Asia, North and South America, Europe, and Oceania; Table 1).

### SNP detection and analysis

SNP analysis was carried out using the RedDog pipeline (https://github.com/katholt/RedDog), which implements a workflow for short read length sequencing analysis (Edwards et al., 2016). Paired-end reads for a total of 422 *B. anthracis* isolates, a single *Bacillus cereus* isolate (chromosomal outgroup), and a single *Bacillus biovar anthracis* isolate (pXO1 and pXO2 plasmid outgroups), were mapped to the *B. anthracis* ‘Ames Ancestor’ reference genome, and its accompanying pXO1 and pXO2 plasmids (GenBank: GCA_000008445.1), using Bowtie2 version 2.2.9 (Langmead & Salzberg, 2012). SNPs were called using SAMTools version 1.8 (Li, 2011). Sites for which conclusive allele calls could not be determined in at least 95% of all isolate sequences at a read depth > 10 were not used, resulting in a final dataset of 356 *B. anthracis* isolates. The resulting SNP dataset then underwent extensive post-processing to (a) remove calls found in regions with large inexact repeats, (b) remove calls within prophage regions of the reference genome identified with PHAST (Zhou et al., 2011), and (c) remove calls from regions that were found to be invariable in all but the outgroup. We also removed calls from regions potentially resulting from recombination, identified by Gubbins (Croucher et al., 2014), or stutter, identified as multiple calls occurring within a 3bp window, or 3 or more calls within a 10bp window. This resulted in a final set of 14,601 chromosomal SNPs in an alignment length of 5,227,419 for the global collection of 356 *B. anthracis* isolates, a final set of 582 SNPs in an alignment length of 181,677 for the accompanying pXO1 plasmid, and a final set of 328 SNPs in an alignment length of 94,830 for the accompanying pXO2 plasmid.

### Phylogenetic and population genomic analysis

We built maximum likelihood (ML) phylogenetic trees from the core genome alignment consisting of 14,601 SNPs (Figure 1a, Figure 2 and Figure 3), from the pXO1 plasmid alignment consisting of 582 SNPs (Figure 1b), and from the pXO2 plasmid alignment consisting of 328 SNPs (Figure 1c) using RAxML version 8.2.4 (Stamatakis, 2014). We applied a generalized time-reversible model and a Gamma distribution to model site specific rate variation. All trees were outgroup rooted, and support for the ML phylogeny was evaluated across 100 bootstrap pseudo-analyses of the alignment for each bacterial component. For visualization, trees were rooted via their outgroups, and arranged in decreasing node order. Outgroups were then removed and all ML trees were visualized and annotated using iTOL version 4.3 (Letunic & Bork, 2016).

**Figure 1.**
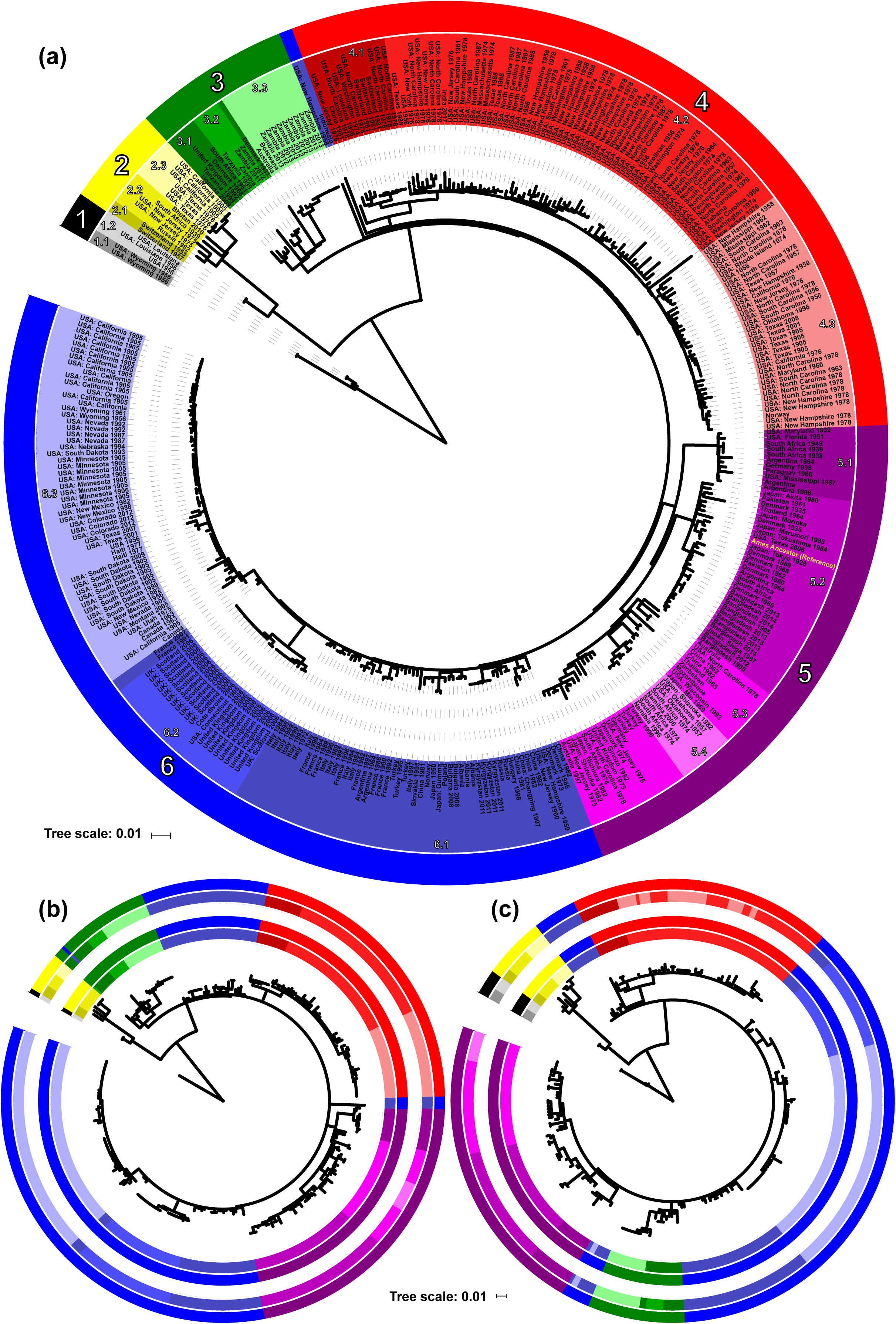
Phylogenetic population structure of B. anthracis based on chromosomal and plasmid wide SNP data. (a) Whole-chromosome tree of 356 global B. anthracis isolates. Primary clusters 1-6 are designated in outer colored ring. Primary clusters are further divided into nested clades indicated with shaded tip labels. The location of the B. anthracis reference genome (accession number GCA_000008445.1) within cluster 5 is labeled in yellow font. (b) Whole-plasmid tree of 283 global B. anthracis pXO1 plasmid isolates. Primary clusters and nested clades based on the chromosomal analysis are designated in outer colored ring. Primary clusters and nested clades based on the plasmid alone are designated in the inner colored ring. (c) Whole-plasmid tree of 268 global B. anthracis pXO2 plasmid isolates. Primary clusters and clades from the chromosomal analysis are designated in outer colored ring. Primary clusters and clades based on the plasmid alone are designated in the inner colored ring.

**Figure 2.**
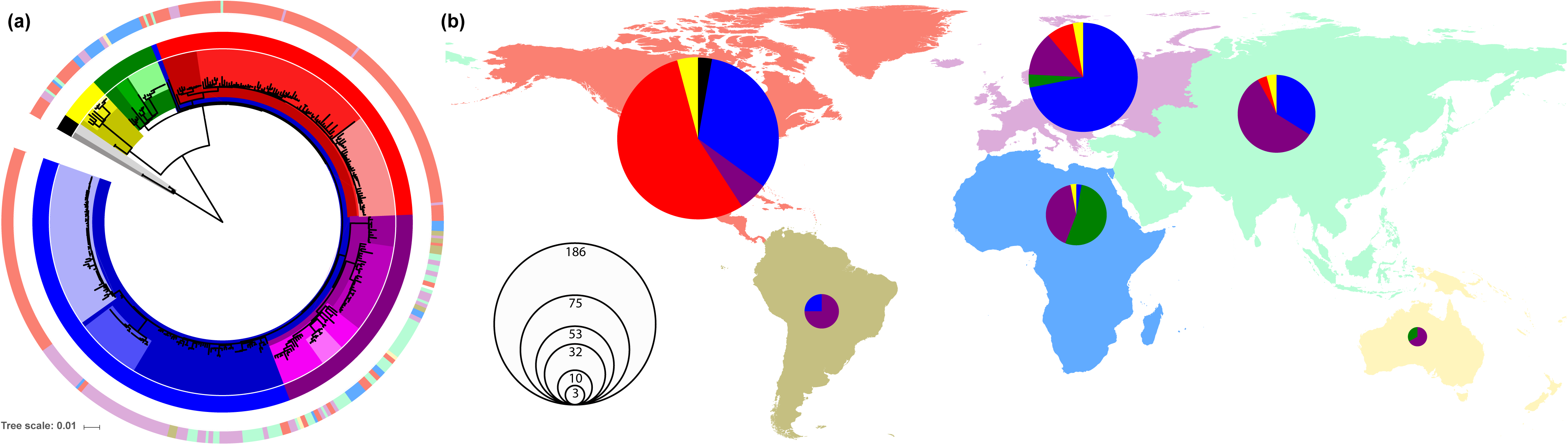
Global population structure of B. anthracis based on core chromosomal SNP architecture. (a) Whole-genome tree of 356 global B. anthracis isolates. Continental collection sites are indicated in the outer colored ring according to the map. The inner ring and shading are colored according to their designation in Figure 1a. (b) Map of the world showing primary cluster diversity of B. anthracis isolates, by continent. North America is colored in red, South America in brown, Europe in purple, Africa in blue, Asia in green, and Oceania in yellow. Pie sizes indicate the number of isolates; slices are colored by primary cluster designation.

**Figure 3.**
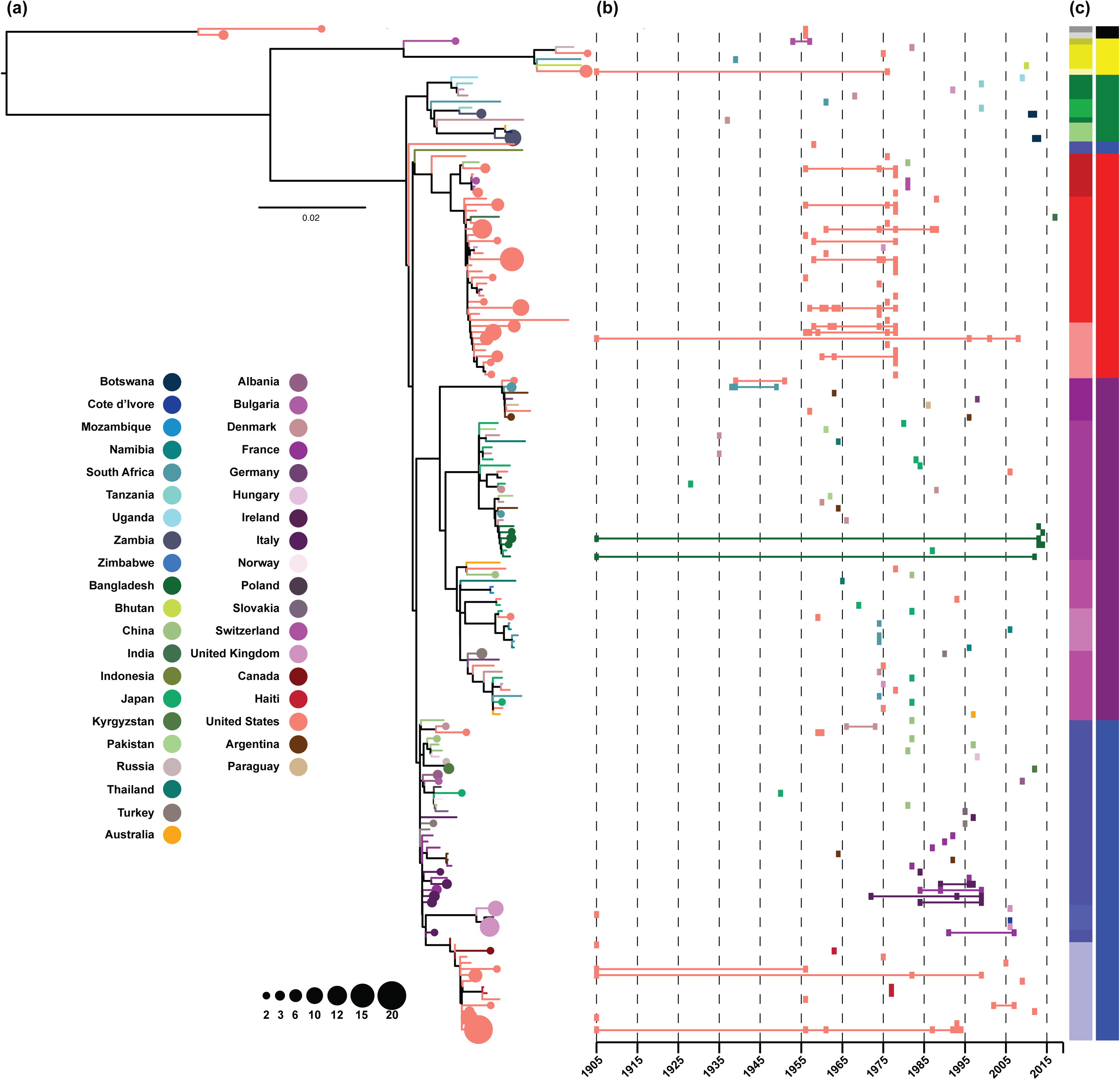
Geographical persistence and diversity of B. anthracis lineages. (a) Maximum likelihood tree for the chromosomal SNP dataset with clades containing branches from a single country collapsed into nodes (circles), sized to indicate the number of isolates in the sub-population and colored by region of isolation. Branches are also colored to indicate the country of origin and correspond to the nodes. (b) Years of isolation for each phylogeographical cluster in the tree. Isolation dates are indicated by rectangles and lines spanning the earliest and latest years of isolation for each cluster are colored to indicate the country of origin. The two adjacent color strips correspond to the clade and primary cluster designations from figures 1 and 2.

To compliment the whole-genome tree building analysis described above, we also applied a population genomic approach to identify distinct clusters across the phylogenetic datasets. For this analysis we employed hierBAPS (Cheng et al., 2013; Tonkin-Hill et al., 2018) which provides a method for hierarchically clustering DNA sequence data to reveal nested population structure. Two nested levels of molecular variation were fitted to the data, and the algorithm was run until it converged at a local optimum for each bacterial component. Clusters identified by hierBAPS were then added to the phylogenetic trees, again employing iTOL to visualize the nested population structure across all *B. anthracis* isolates (Figure 1, Figure 2, and Figure 3). To understand how geographical and temporal differences related to tree topology, branches that included multiple isolates from the same country were manually collapsed using Figtree version 1.4.3 (Rambaut, 2009), and annotated with sample size and collection dates using Adobe Illustrator CC Adobe Inc. 2019 (Figure 3).

### SNP-based genotyping framework

We distinguished a minimum set of 19 canonical chromosomal SNPs that can be used to rapidly categorize *B. anthracis* isolates into the 6 primary clusters and 18 nested clades described above (S1 Table). A multi-variant call file, generated by mapping Illumina reads to the Ames Ancestor reference genome (accession AE017334.2) was used to assign genotypes for each unique read set using the program Gingr (Treangen et al., 2014)^28^. Gingr is an interactive tool for exploring large-scale phylogenies in tandem with their corresponding multi-alignments, and was used to identify cluster-, and clade-defining SNP alleles across the chromosomal alignment.

## RESULTS

### Defining a classification framework using the *B. anthracis* core genome

We used WGS data from 356 globally representative *B. anthracis* isolates to establish a phylogenetically informative framework based on core population genomic SNP architecture. Summary information on isolates is presented in Table 1 and detailed information for isolates are available in S1 Dataset. Phylogenetic tree topology was integrated into a population genomic framework, allowing for the identification of 6 primary *B. anthracis* clusters that could be further sub-divided into 18 nested clades (Figure 1a). Each major cluster exhibited 100% bootstrap support and was delineated by > 20 SNPs. The mean pairwise difference between isolates at the cluster level was 273 SNPs, while the mean pairwise distance between isolates at the clade level was 124 SNPs. Clusters and clades were named using a hierarchical system, such as those used for *M. tuberculosis* and *Salmonella typhi* (Coll et al., 2014; Wong et al., 2016). In this naming system, each primary cluster is designated a number, such as 1, and nested clades within that cluster are indicated by decimals, such as 1.1 and 1.2.

After defining a genotyping framework, we determined a minimum set of 19 SNPs that can be used to accurately designate *B. anthracis* into the 6 defined primary clusters and 18 subsequent clades for classification purposes. We chose SNPs from protein-coding sequences that exhibited low nucleotide diversity across the dataset, with no identifiable mutations potentially leading to inactivation. We also developed a python script to easily classify *B. anthracis* draft genomes (mapped to the reference using parsnp; Treangen et al., 2014) or mapped reads (in either BAM of VCF format) for other researchers seeking to utilize this system (S1 File). Additional information regarding genotyping SNPs can be found in S1 Table. We have also made the RedDog output folder associated with this study available for researchers seeking to append their WGS data in the context of the full analyses. This classification framework provides greatly improved discriminatory power when compared to the older classification system (S2 Fig), is phylogenetically descriptive in form, and the hierarchical nomenclature used to define both primary clusters and nested clades is fundamentally informative by design.

### Comparing *B. anthracis* core genome and plasmid structure

We examined differences in the genomic structure of *B. anthracis* plasmids compared to the core genome, to understand inheritance patterns, and identify how plasmid evolution may differ from that exhibited in the core phylogeny. Plasmid tree building followed the same methodology as the core genome, and the resulting datasets were again classified using a population genomic framework to elucidate two levels of nested structure across both the pXO1 and pXO2 plasmids (Figure 1b and 1c, see methods). For both plasmids a subset of samples failed to meet read depth criteria across the replicon due to absence or low count number, resulting in a total of 283 samples used to build the pXO1 tree, and 268 samples used to build the pXO2 tree. Samples that failed to meet the read depth criteria for each plasmid are listed in S3 Table. Tree structure across both plasmids was generally similar to that of the core genome (Figure 1). The hierBAPS package assigned individual isolates to the same major clusters as for the core phylogeny for both the pXO1 and pXO2 plasmids, with the exception of a single sample from Poland in the pXO1 tree. This sample was assigned to primary cluster 3, rather than its chromosomal designation to primary cluster 6. These results suggest that the plasmids are primarily inherited vertically, and have coevolved with the bacterial chromosome. Ambiguities at the clade level were minimal, and usually resulted from a single decrease in the number of identifiable clades within a major cluster, with the exception of several outliers from cluster 6, and the arrangement of cluster 3 in plasmid pXO2 along the phylogenetic backbone, likely due to decreased resolution given the small size (∼ 90,000 bp) and limited number of SNPs detected across the pXO2 replicon (328).

### Geographical and temporal distribution of *B. anthracis*

We also examined the geographical and temporal distribution of *B. anthracis* isolates based on our global genotyping framework. When examining the dataset with regards to continental distribution, most primary clusters spanned multiple continents, putatively reflecting the spread of *B. anthracis* relative to both ancient dissemination and contemporary transit in terms of agricultural trade practices (Figure 2). The only exception to this is primary cluster 1 (black; Figure 2), which contains a limited number of representative isolates (*n* = 5) and are all from North America (consistent with the previously described major lineage C). When examining all other primary clusters, some clusters were more heavily weighted with samples from certain regions than others. The majority of isolates representative of primary cluster 2 (yellow; Figure 2) were from North America (*n* = 8; 61%), but isolates in this cluster were also found in Africa (*n* = 1; 8%), Europe (*n* = 2; 15%), and Asia (*n* = 2; 16%) (consistent with the previously described major lineage B). Primary clusters 3 through 6 (green, red, purple, blue; Figure 2) were all consistent with the previously described major lineage A. Primary cluster 3 was dominated by isolates from Africa (*n* = 17; 81%), but also contained isolates from Europe (*n* = 3; 14%) and Oceania (*n* = 1; 5%). Primary cluster 4, which makes up a large portion of the isolates examined in this study, was dominated by isolates from of North America (*n* = 102; 93%), but also contained several isolates from Europe (*n* = 6; 5%) and Asia (*n* = 2; 2%). Primary cluster 5 represents the most geographically, and genetically diverse primary cluster examined (mean number of pairwise SNPs between clades > 133). The majority of isolates from this cluster are from Asia (*n* = 31; 43%), but this cluster also contained multiple representatives from all other continental regions examined in this study. Primary cluster 6 is the largest cluster examined in this study, and is dominated by isolates from North America (*n* = 62; 46%) and Europe (*n* = 54; 40%), but also contains several isolates from Asia (*n* = 18; 14%), and South America (*n* = 2; 1%). Clade 6.3 from primary cluster 6 is made up entirely of isolates from North America (*n* = 56) and is the largest single clade in the dataset. Clade 6.2 contains a single sample from Africa, the only such sample found across 135 isolates in primary cluster 6. Ancestral isolates from primary clusters 1 and 2 which result from deep branching originating in close proximity to the root of the *B. anthracis* whole genome tree, were rare in the collection (*n* = 18) and mostly found in North America (*n* = 13; 72%). Nevertheless, it should be noted that this dataset is heavily weighted by isolates collected in North America, accounting for more than half of the isolates examined in study (*n* = 186; 52%).

Finally, the whole genome tree was examined at the level of individual countries to assess temporal patterns in genomic structure across the genotyping framework, for samples for which dates were available. In this analysis, branches within a designated clade specific to a country that contained more than one isolate were collapsed and plotted across years of isolation for each phylogeographical branch in the tree (Figure 3). The 5 samples representative of primary cluster 1 were all isolated in the United States in the same year; 1956, while samples representative of primary cluster 3 spanned the time period between 1937 and 2013. All other primary clusters contained at least one sample isolated as early as 1905, suggesting the spread of many *B. anthracis* lineages predates isolate collection for this study. Isolates representative of primary clusters 2 and 3 were both geographically and temporally diverse, spanning multiple decades and oceanic barriers. The majority of isolation dates for primary cluster 4 fell between 1955 and 1980, and were relegated to the United States (*n* = 102) with the exception of one sample from the United Kingdom. Other exceptions that fell outside of this date range included several samples from Switzerland (*n* = 3), and a sample from China isolated in 1981, as well as single sample from India isolated in 2017. One collapsed branch in this cluster exhibited samples ranging in isolation date from 1905 to 2008, suggesting that this lineage has likely persisted in the United States for over a century. Isolation dates and countries of origin for primary cluster 5 varied widely with the majority of samples exhibiting isolation dates between 1955 and 2015. Some exceptions included several samples from Denmark isolated in 1935 (*n* = 2), samples from South Africa isolated between 1938 and 1949 (*n* = 3), and a sample from Japan isolated in 1927. Bangladesh exhibited several branch-specific isolates with dates that ranged between 1905 and 2014, suggesting lineages from this clade have similarly persisted in this region for over a century. Samples representative of primary cluster 6 change in terms of both origin and isolation date as one follows the backbone of the phylogenetic tree downward. The upper portion of the cluster is geographically diverse with the majority of samples exhibiting isolation dates that fall between 1965 and 2000. Some exceptions to this rule include a sample from Japan isolated in 1950, and two samples from the United States isolated in 1958 and 1959. As one moves downwards along the tree, primary cluster 6 becomes dominated with samples collected in Europe, primarily isolated between 1970 and 2000 (*n* = 54). The final portion of primary cluster 6, defined by clade 6.3 is exclusive to the United States and Canada (*n* = 62), exhibiting isolation dates that range well over a century, suggesting that this lineage may be uniquely suited to persistence in this region. For the most part, isolation dates and countries of origin vary widely across primary clusters, confirming an ancient global expansion of *B. anthracis* across oceanic barriers and landmasses (Van Ert et al., 2007a).

## DISCUSSION

The data presented in this study demonstrate that the global *B. anthracis* population consists of 6 distinct primary clusters, and 18 clades that are geographically structured in most cases, with the exception of primary cluster 5 which exhibited both high genomic and geographic diversity. Nevertheless, many of the regions examined harbor diverse anomalies likely established before isolate collection began, reinforcing the inferred ancient global dissemination and persistence of this bacterium. We also demonstrate that the *B. anthracis* plasmids exhibit similar population structure to the core genome, suggesting that plasmid inheritance is vertical in the majority of cases. An exception to this rule was a single isolate collected in Poland, whose pXO1 plasmid structure was consistent with primary cluster 3, despite its chromosomal designation to primary cluster 6.

The primary clusters and nested clades for the *B. anthracis* chromosome can be identified through a simple genotyping framework consisting of only 19 SNPs. Notably, we also show that this genotyping framework is highly informative and can be readily inferred from draft genome sequences or mapped reads without the need for laborious whole-genome comparisons or other computationally expensive analyses. These properties suggest that this framework may act as a powerful tool for researchers and institutions looking to develop future studies around *B. anthracis* population genomic structure. In addition, given the level of stability in terms of minimal genomic variation and sparse evidence of recombination found in the *B. anthracis* genome (Harrell et al., 1995; Zwick et al., 2005), the genotyping system presented here can be expected to maintain its relevancy for many years to come.

When comparing the classification framework with the previous system, major differences in resolution are easily discernable, ultimately leading to higher specificity in sample designation for the revised approach. Primary cluster 1 (consistent with the previously described major lineage C; S2 Fig) was unclassifiable using the older canSNPer-based scheme, whereas the new framework allows for both primary cluster designation, as well as clade designation within this group. When examining primary cluster 2 (consistent with the previously described major lineage B; S2 Fig), we see that the previous system allowed for only two sub-groups within our dataset, where as a test of population genomic structure detected and designated 3. In addition, when viewing the new framework in the context of previously described major lineage A, we were able to distinguish 4 major clusters, and 10 nested clades, whereas the previous scheme allowed for only 4 subgroups and 3 additional sub-lineages based on rare alleles in our dataset. Lastly, the older genotyping scheme exhibited an instance of misclassification for a single sample falling on the edge of A.Br.009 (classified as A.Br.004; S2 Fig), suggesting homoplasy and indicating that further misclassification upon expansion of the dataset has the potential to produce erroneous results. This updated genotyping scheme therefore allows for a far more robust system for tracking, distinguishing and tracing the evolutionary origins of *B. anthracis* based on global genomic structure.

As a result of the geographic clustering exhibited at the clade level, the incorporation of additional *B. anthracis* isolates may lead to an even more robust predictive genotyping scheme, helping to identify the geographic origins of regional outbreaks at finer scales. For example, two samples that fall between primary clusters 3 and 4, from the United States and Indonesia were designated to clade 6.1 despite being highly differentiated from each other and all other isolates examined this study (mean number of pairwise SNP differences from all other isolates > 430, and > 567, respectively). With the addition of genetically similar isolates, these lone branches may eventually form their own clades, or even primary clusters, and could help to expand the current framework. A refined and accurate genotyping scheme has important wildlife conservation, livestock management, and human health implications for *B. anthracis* surveillance and management; however, continued updates to the genotyping framework will be essential to ensuring the practicality of this type of monitoring.

Resolving the geo-evolutionary origins of *B. anthracis* based on contemporary samples remains problematic, as the geographical spread of *B. anthracis* likely pre-dates sample collection by many thousands of years (Van Ness, 1971; Kenefic et al., 2009; Van Ert et al., 2007a). Past work on the global genomic structure of *B. anthracis* has attributed the level of global diversity found in this pathogen to independent domestication, domesticate expansion events, and more recent human activities such as transcontinental commerce, and industrialization (Tatem et al., 2006; Wyres et al., 2014). Work that has specifically examined the distribution and spread of *B. anthracis* in Western North America has suggested that the lineage specific to this region (clade 6.3; formerly known A.Br.009) may have arrived via the Bering Land Bridge (Kenefic et al., 2009). When examining continental diversity in terms of phylogeographic structure, the results presented here contradict those findings suggesting this lineage is consistent with European origins. In addition, the results of this study suggest that some clades may have potentially evolved to fill ecological niches based on sub-groupings that appear to be biogeographically suited to specific regions. With the expansion of this genotyping system a more robust understanding of *B. anthracis* evolutionary specificity may be possible, especially with the inclusion of isolates from the many regions that are currently under-represented in this dataset.

Over the past several decades many public health institutions and wildlife disease laboratories have replaced PCR-based methods with WGS, given the falling price of sample throughput and high-resolution data available from such technology (Wyres et al., 2014). This provides a fantastic opportunity to utilize and expand the global framework presented here, with potential benefits for both local and broader regional agencies seeking to understand both outbreak origins and recent introductions of this bacterium. It is our hope that the genomic framework presented in this study may offer insight as to how WGS technology can be used to better understand global distributions and answer complex questions regarding pathogen evolution and dissemination across the globe.

## DATA ACCESSIBILITY

Raw sequence data used in this study are available through the NCBI database and accession numbers are provided in S1 Dataset. The RedDog pipeline is available at https://github.com/katholt/RedDog. The RedDog output folder created for this study can be accessed through the Dryad database, and may be used to expand upon the SNP dataset by merging new runs with the existing output via RedDog. In addition, a Python script for the new genotyping framework has also been made available for rapid primary cluster and clade designation using draft genomes mapped to the reference via parsnp, or mapped reads in the form of BAM or VCF files (S1 File). Data for this study are available at: to be completed after manuscript is accepted for publication

## Acknowledgments

This work is supported by the NSF Division of Environmental Biology, grant number 1816161 (W.C.T.) and start-up funds from the University at Albany (W.C.T.).

