## Supplementary material for "A classification framework for *Bacillus anthracis* defined by global genomic structure": S2 Fig

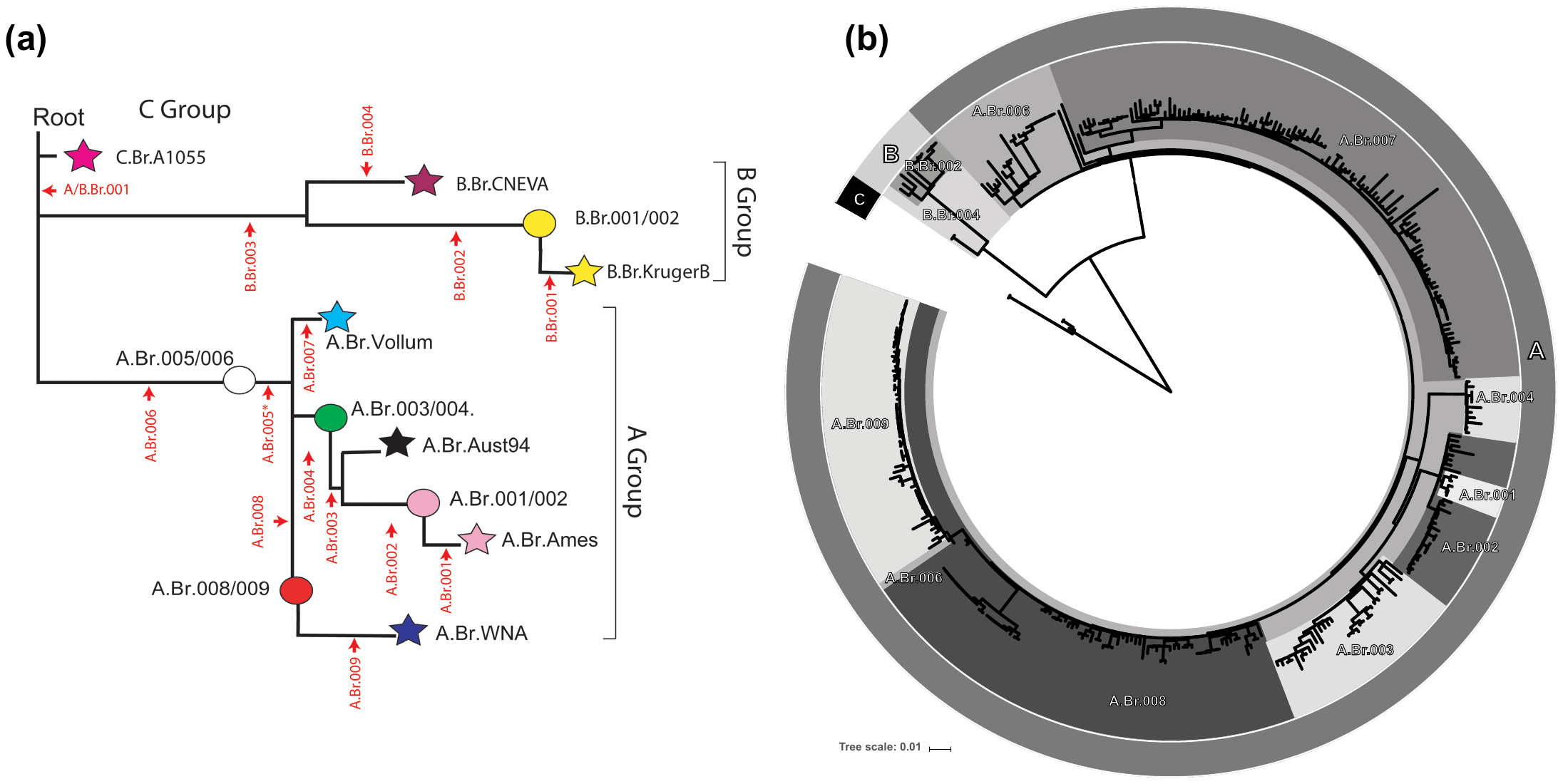
**S1 Fig. Chromosomal phylogeny for the *B. anthracis* global dataset in relation to the Van Ert *et al.* classification scheme.** (a) The relationship between canSNPs, sub-lineages and/or sub-groups from Van Ert et al., 2007: The stars in this dendrogram represent specific lineages that are defined by one of the seven sequenced genomes of B. anthracis. The circles represent branch points along the lineages that contain specific subgroups of isolates. These sub-groups are named after the canSNPs that flank these positions. Indicated in red are the positions and names for each of the canSNPs (b) Whole-chromosome tree of 356 global *B. anthracis* isolates shaded and labeled using the genotyping scheme shown in a*.* De novo assemblies were performed on paired raw read files using SPAdes, and classified with the CanSNPer package. All tree annotation was carried out using iTOL.

**S1 Table**

| Group | N | In | Out | Position | Locus | Gene |
| --- | --- | --- | --- | --- | --- | --- |
| 1.1 | 2 | A | T | 1771938 | GBAA_1887 | enterotoxin A |
| 1.2^a^ | 3 | T | C | 1201909 | GBAA_1246 | sodium/proline symporter family protein |
| 2.1 | 2 | A | G | 2212885 | GBAA_2372 | dhbf |
| 2.2^b^ | 5 | G | A | 2132895 | GBAA_2284 | hypothetical protein |
| 2.3^c^ | 6 | A | G | 2019941 | GBAA_2169 | transcriptional regulator, TetR family |
| 3.1 | 6 | C | A | 3086718 | GBAA_3355 | alo |
| 3.2^d^ | 4 | C | T | 3366432 | GBAA_3657 | parE |
| 3.3^d^ | 11 | T | G | 3359543 | GBAA_3652 | gene |
| 4.1 | 12 | C | T | 810017 | GBAA_0794 | hypothetical protein |
| 4.2^e^ | 63 | T | C | 837700 | GBAA_0825 | conserved hypothetical protein |
| 4.3^f^ | 35 | G | A | 1271875 | GBAA_1327 | phaP protein |
| 5.1 | 11 | T | C | 857338 | GBAA_0848 | conserved hypothetical protein |
| 5.2^g^ | 29 | C | A | 4499321 | GBAA_4954 | hypothetical protein |
| 5.3^g^ | 24 | T | C | 569295 | GBAA_0561 | citrate transporter, CitM family |
| 5.4^h^ | 8 | T | C | 565120 | GBAA_0558 | methyl-accepting chemotaxis protein |
| 6.1 | 57 | T | C | 2332200 | GBAA_2510 | csbX |
| 6.2^i^ | 22 | T | C | 2387683 | GBAA_2566 | acetyltransferase, GNAT family |
| 6.3^i^ | 56 | A | G | 1776654 | GBAA_1891 | putative deoxyribonucleoside regulator DeoR |

**S1 Table. Canonical SNPs for classifying *B. anthracis* isolates.** Key: Group = Primary cluster/Clade; N = number of *B. anthracis* isolates in the Group; In = allele within the group; Out = allele outside of group; ^a^ nested in 1.1; ^b^ nested in 2.1; ^c^ nested in 2.2; ^d^ nested in 3.1; ^e^ nested in 4.1; ^f^ nested in 4.2; ^g^ nested in 5.1; ^h^ nested in 5.3; ^i^ nested in 6.1.

**S2 Table**

| Accession | Origin | Isolation date | Failed bacterial component |
| --- | --- | --- | --- |
| SRR5810989 | Argentina | 1964 | pXO2 |
| SRR5811192 | Argentina | 1964 | pXO1, pXO2 |
| SRR2968170 | Botswana | - | pXO2 |
| SRR2339291 | USA: California | 1976 | Chromosome, pXO1, pXO2 |
| SRR2339617 | USA: California | 1976 | Chromosome, pXO1, pXO2 |
| SRR2340465 | USA: California | 1976 | Chromosome, pXO1, pXO2 |
| SRR5811182 | USA: California | 1976 | pXO2 |
| SRR5811208 | USA: California | 1976 | pXO1, pXO2 |
| SRR5811191 | Canada | 1963 | pXO1, pXO2 |
| SRR5811247 | Canada | 1963 | pXO1 |
| SRR2157175 | USA: Florida | 1951 | pXO2 |
| SRR2339516 | Haiti | 1976 | Chromosome, pXO1, pXO2 |
| SRR2339619 | Haiti | 1976 | Chromosome, pXO1, pXO2 |
| SRR2340858 | Haiti | 1977 | Chromosome, pXO1, pXO2 |
| SRR5811210 | Haiti | 1977 | pXO1 |
| SRR2340466 | Haiti | 1981 | Chromosome, pXO1, pXO2 |
| SRR2340467 | USA: Iowa | 1979 | Chromosome, pXO1, pXO2 |
| SRR3418280 | Kyrgyzstan | 2011 | pXO2 |
| SRR3418281 | Kyrgyzstan | 2011 | pXO2 |
| SRR3418282 | Kyrgyzstan | 2011 | pXO2 |
| SRR3418292 | Kyrgyzstan | 2011 | pXO2 |
| SRR5811162 | USA: Louisiana | 1956 | pXO1, pXO2 |
| SRR5947105 | USA: Louisiana | 1956 | pXO1 |
| SRR2145877 | USA: Louisiana | - | pXO1 |
| SRR2145878 | USA: Louisiana | - | Chromosome, pXO1, pXO2 |
| SRR5811216 | USA: Maryland | 1960 | pXO1, pXO2 |
| SRR5810964 | USA: Massachusetts | 1974 | pXO1, pXO2 |
| SRR5810965 | USA: Massachusetts | 1974 | pXO2 |
| SRR5811041 | USA: Massachusetts | 1974 | pXO1, pXO2 |
| SRR2340910 | USA: Massachusetts | 1976 | Chromosome, pXO1, pXO2 |
| SRR5810983 | USA: Mississippi | 1957 | pXO1 |
| SRR5811070 | USA: Mississippi | 1962 | pXO1 |
| SRR5811168 | USA: Mississippi | 1962 | pXO1, pXO2 |
| SRR5810951 | USA: New Hampshire | 1958 | pXO1 |
| SRR5810952 | USA: New Hampshire | 1958 | pXO1, pXO2 |
| SRR5810953 | USA: New Hampshire | 1958 | pXO2 |
| SRR5810954 | USA: New Hampshire | 1958 | pXO1, pXO2 |
| SRR5810991 | USA: New Hampshire | 1958 | pXO2 |
| SRR5811248 | USA: New Hampshire | 1958 | pXO1, pXO2 |
| SRR5811099 | USA: New Hampshire | 1959 | pXO2 |
| SRR5811100 | USA: New Hampshire | 1959 | pXO2 |
| SRR2339640 | USA: New Hampshire | 1978 | Chromosome, pXO1, pXO2 |
| SRR2339645 | USA: New Hampshire | 1978 | Chromosome, pXO1, pXO2 |
| SRR2340010 | USA: New Hampshire | 1978 | Chromosome, pXO1, pXO2 |
| SRR2340073 | USA: New Hampshire | 1978 | Chromosome, pXO1, pXO2 |
| SRR2340171 | USA: New Hampshire | 1978 | Chromosome, pXO1, pXO2 |
| SRR2340230 | USA: New Hampshire | 1978 | Chromosome, pXO1, pXO2 |
| SRR2340486 | USA: New Hampshire | 1978 | Chromosome, pXO1, pXO2 |
| SRR2340857 | USA: New Hampshire | 1978 | Chromosome, pXO1, pXO2 |
| SRR5810966 | USA: New Hampshire | 1978 | pXO1 |
| SRR5810967 | USA: New Hampshire | 1978 | pXO2 |
| SRR5810968 | USA: New Hampshire | 1978 | pXO1, pXO2 |
| SRR5810969 | USA: New Hampshire | 1978 | pXO2 |
| SRR5810970 | USA: New Hampshire | 1978 | pXO1, pXO2 |
| SRR5810971 | USA: New Hampshire | 1978 | pXO1, pXO2 |
| SRR5810972 | USA: New Hampshire | 1978 | pXO2 |
| SRR5811006 | USA: New Hampshire | 1978 | pXO1, pXO2 |
| SRR5811147 | USA: New Hampshire | 1978 | pXO2 |
| SRR5811230 | USA: New Hampshire | 1978 | pXO2 |
| SRR5811234 | USA: New Hampshire | 1978 | pXO1, pXO2 |
| SRR2340305 | USA: New Jersey | 1975 | Chromosome, pXO1, pXO2 |
| SRR5810962 | USA: New Jersey | 1960 | pXO1, pXO2 |
| SRR2340252 | USA: New Jersey | 1975 | pXO1 |
| SRR5811079 | USA: New Jersey | 1975 | pXO1, pXO2 |
| SRR5811124 | USA: New Jersey | 1975 | pXO1, pXO2 |
| SRR5811125 | USA: New Jersey | 1975 | pXO1 |
| SRR5811126 | USA: New Jersey | 1975 | Chromosome, pXO1, pXO2 |
| SRR2340461 | USA: New Jersey | 1976 | Chromosome, pXO1, pXO2 |
| SRR2340463 | USA: New Jersey | 1976 | Chromosome, pXO1, pXO2 |
| SRR2340464 | USA: New Jersey | 1976 | Chromosome, pXO1, pXO2 |
| SRR2895370 | USA: New Jersey | 1976 | Chromosome, pXO1, pXO2 |
| SRR5811181 | USA: New Jersey | 1976 | pXO1, pXO2 |
| SRR5811184 | USA: New Jersey | 1976 | pXO1, pXO2 |
| SRR5811185 | USA: New Jersey | 1976 | pXO1, pXO2 |
| SRR5811225 | USA: New Jersey | 1976 | pXO2 |
| SRR5811209 | USA: New York | 1976 | pXO1 |
| SRR5811005 | USA: North Carolina | 1957 | pXO1, pXO2 |
| SRR5811061 | USA: North Carolina | 1957 | pXO1, pXO2 |
| SRR5811050 | USA: North Carolina | 1961 | pXO1 |
| SRR2339241 | USA: North Carolina | 1978 | Chromosome, pXO1, pXO2 |
| SRR2340468 | USA: North Carolina | 1978 | Chromosome, pXO1, pXO2 |
| SRR2340469 | USA: North Carolina | 1978 | Chromosome, pXO1, pXO2 |
| SRR2340470 | USA: North Carolina | 1978 | Chromosome, pXO1, pXO2 |
| SRR2340471 | USA: North Carolina | 1978 | Chromosome, pXO1, pXO2 |
| SRR2340474 | USA: North Carolina | 1978 | Chromosome, pXO1, pXO2 |
| SRR2340475 | USA: North Carolina | 1978 | Chromosome, pXO1, pXO2 |
| SRR2340476 | USA: North Carolina | 1978 | Chromosome, pXO1, pXO2 |
| SRR2340477 | USA: North Carolina | 1978 | Chromosome, pXO1, pXO2 |
| SRR2340478 | USA: North Carolina | 1978 | Chromosome, pXO1, pXO2 |
| SRR2340480 | USA: North Carolina | 1978 | Chromosome, pXO1, pXO2 |
| SRR2340481 | USA: North Carolina | 1978 | Chromosome, pXO1, pXO2 |
| SRR2340482 | USA: North Carolina | 1978 | Chromosome, pXO1, pXO2 |
| SRR2340483 | USA: North Carolina | 1978 | Chromosome, pXO1, pXO2 |
| SRR2340485 | USA: North Carolina | 1978 | Chromosome, pXO1, pXO2 |
| SRR2340487 | USA: North Carolina | 1978 | Chromosome, pXO1, pXO2 |
| SRR2340488 | USA: North Carolina | 1978 | Chromosome, pXO1, pXO2 |
| SRR2340490 | USA: North Carolina | 1978 | Chromosome, pXO1, pXO2 |
| SRR2340492 | USA: North Carolina | 1978 | Chromosome, pXO1, pXO2 |
| SRR2340505 | USA: North Carolina | 1978 | Chromosome, pXO1, pXO2 |
| SRR2340670 | USA: North Carolina | 1978 | Chromosome, pXO1, pXO2 |
| SRR2340697 | USA: North Carolina | 1978 | Chromosome, pXO1, pXO2 |
| SRR2340730 | USA: North Carolina | 1978 | Chromosome, pXO1, pXO2 |
| SRR2340969 | USA: North Carolina | 1978 | Chromosome, pXO1, pXO2 |
| SRR2340986 | USA: North Carolina | 1978 | Chromosome, pXO1, pXO2 |
| SRR5811016 | USA: North Carolina | 1978 | pXO1, pXO2 |
| SRR5811017 | USA: North Carolina | 1978 | pXO1 |
| SRR5811019 | USA: North Carolina | 1978 | pXO1, pXO2 |
| SRR5811020 | USA: North Carolina | 1978 | pXO1 |
| SRR5811021 | USA: North Carolina | 1978 | pXO1 |
| SRR5811121 | USA: North Carolina | 1978 | pXO1 |
| SRR5811122 | USA: North Carolina | 1978 | pXO1 |
| SRR5811128 | USA: North Carolina | 1978 | pXO1, pXO2 |
| SRR5811129 | USA: North Carolina | 1978 | pXO2 |
| SRR5811178 | USA: North Carolina | 1978 | pXO1, pXO2 |
| SRR5811205 | USA: North Carolina | 1978 | pXO1 |
| SRR5811228 | USA: North Carolina | 1978 | pXO1, pXO2 |
| SRR5811046 | USA: North Carolina | 1987 | pXO2 |
| SRR5811049 | USA: North Carolina | 1987 | pXO2 |
| SRR2339961 | USA: North Carolina | 1988 | Chromosome, pXO1, pXO2 |
| SRR5811048 | USA: North Carolina | 1988 | pXO1 |
| DRR128188 | Japan: Okayama |  | Chromosome, pXO2 |
| SRR5811167 | Pakistan | 1961 | pXO2 |
| SRR2339632 | Paraguay | - | Chromosome, pXO1, pXO2 |
| SRR2339823 | USA: Pennsylvania | 1974 | Chromosome, pXO1, pXO2 |
| SRR5811083 | USA: Pennsylvania | 1974 | pXO1, pXO2 |
| SRR2339757 | USA: Rhode Island | 1974 | Chromosome, pXO1, pXO2 |
| SRR2339940 | USA: Rhode Island | 1974 | Chromosome, pXO1, pXO2 |
| SRR5811213 | USA: Rhode Island | 1974 | pXO1 |
| SRR2968141 | Russia | 1979 | Chromosome |
| SRR2968143 | Russia | 1979 | Chromosome |
| SRR2968198 | Russia | 1979 | Chromosome, pXO1, pXO2 |
| SRR2968216 | Russia | 1979 | Chromosome, pXO1, pXO2 |
| SRR2968168 | Russia | - | pXO2 |
| SRR2968169 | Russia | - | pXO2 |
| SRR5811076 | South Africa | 1961 | pXO2 |
| SRR2339898 | South Africa | 1974 | Chromosome, pXO1, pXO2 |
| SRR5811080 | South Africa | 1974 | pXO2 |
| SRR5811082 | South Africa | 1974 | pXO1, pXO2 |
| SRR5811085 | South Africa | 1974 | pXO1 |
| SRR5811215 | South Africa | 1974 | pXO1 |
| SRR1739968 | South Africa | - | pXO2 |
| SRR5811037 | South Africa | - | pXO2 |
| SRR5810963 | USA: South Carolina | 1956 | pXO1, pXO2 |
| SRR5811003 | USA: South Carolina | 1956 | pXO1, pXO2 |
| SRR2339626 | USA: South Carolina | 1960 | Chromosome, pXO1, pXO2 |
| SRR5811063 | USA: South Carolina | 1960 | pXO1, pXO2 |
| SRR5810959 | USA: South Carolina | 1961 | pXO1, pXO2 |
| SRR5811004 | USA: South Carolina | 1961 | pXO1, pXO2 |
| SRR5810997 | USA: South Carolina | 1963 | pXO1, pXO2 |
| SRR5811000 | USA: South Carolina | 1963 | pXO2 |
| SRR5811007 | USA: South Carolina | 1963 | pXO1, pXO2 |
| SRR5810998 | USA: South Carolina | 1964 | pXO1, pXO2 |
| SRR5811069 | USA: Texas | 1976 | pXO1, pXO2 |
| SRR5811166 | USA: Texas | 1976 | pXO1 |
| SRR5811218 | USA: Texas | 1976 | pXO1 |
| SRR2339631 | USA: Texas | 1988 | Chromosome, pXO1, pXO2 |
| SRR5811145 | USA: Texas | 1988 | pXO2 |
| SRR5811146 | USA: Texas | 1988 | pXO2 |
| SRR5811219 | Thailand | 1964 | pXO1, pXO2 |
| SRR5811187 | Thailand | 1965 | pXO1 |
| SRR2968136 | Turkey | - | pXO2 |
| SRR2968144 | Turkey | - | pXO1 |
| SRR2968193 | Turkey | - | pXO2 |
| SRR2340304 | United Kingdom | 1975 | Chromosome, pXO1, pXO2 |
| SRR5811120 | United Kingdom | 1975 | pXO2 |
| SRR5811123 | United Kingdom | 1975 | pXO1, pXO2 |
| SRR2094267 | United Kingdom | 2006 | pXO2 |
| SRR2094270 | United Kingdom | 2006 | pXO2 |
| SRR2094273 | United Kingdom | 2006 | pXO2 |
| SRR5811034 | United States | 1956 | pXO2 |
| SRR5811101 | United States | 1956 | pXO1, pXO2 |
| SRR5811103 | United States | 1956 | pXO1, pXO2 |
| SRR5811104 | United States | 1956 | pXO1, pXO2 |
| SRR2340460 | USA: Utah | 1975 | Chromosome, pXO1, pXO2 |
| SRR5811084 | USA: Utah | 1975 | pXO1, pXO2 |
| SRR2339728 | USA: Washington | 1974 | Chromosome, pXO1, pXO2 |
| SRR5811040 | USA: Washington | 1974 | pXO1, pXO2 |
| SRR5811078 | USA: Washington | 1974 | pXO2 |
| SRR5811081 | USA: Washington | 1974 | pXO1, pXO2 |
| SRR5811086 | USA: Washington | 1974 | pXO2 |
| SRR5811190 | USA: Washington | 1974 | pXO2 |
| SRR1739967 | USA: Wyoming | 1956 | pXO1 |
| SRR5811161 | USA: Wyoming | 1956 | pXO1, pXO2 |
| SRR5811214 | USA: Wyoming | 1956 | pXO1 |
| SRR5811096 | USA: Wyoming | 1961 | pXO1 |
| DRR014738 | Zambia | 2013 | Chromosome, pXO1, pXO2 |

**S2 Table. Isolate information for individuals that failed to meet coverage and read depth criteria**. Isolates listed below failed to meet coverage and read depth criteria (see methods) when mapped to the Ames Ancestor reference genome (accession number GCA_000008445.1; Chromosome: AE017334, pXO1 replicon: AE017336, pXO2 replicon: AE017335).

**S1 Dataset. Metadata for all isolates included in this study.** This excel file includes strain information, NCBI accession numbers, years of isolation (if known), continent, country/region of origin, isolation source, genotype classification and submission source information for all isolates.

**S1 File. Python script for rapid *B. anthracis* genotyping.** This script can be used for rapid primary cluster and clade designation using *B. anthracis* draft genomes (mapped to a reference) or mapped raw read data in the form of BAM or VCF files.
