## Supplementary material for "A classification framework for *Bacillus anthracis* defined by global genomic structure": S1 File

#!/usr/bin/env python
#
# Input BAM (recommended) or VCF (if highly trusted SNP data) relative to Ames Ancestor (AE017334) and assign B.anthracis genotype codes.
#
# Authors - Originally written for S.typhi by Kat Holt and Zoe Dyson
# Modified for B. anthracis by Spencer Bruce and Nick Schiraldi
#
# Original documentation - https://github.com/katholt/genotyphi
#
# Dependencies:
#	 SAMtools (v1.2) and bcftools (v1.2) are required to genotype from BAMs.
#
# Last modified - Apr 4th, 2019
#

from argparse import (ArgumentParser, FileType)
import os, sys, re, collections, operator
import gzip
import logging
import time
import datetime
from subprocess import call, check_output, CalledProcessError, STDOUT


def parse_args():
	"Parse the input arguments, use '-h' for help"
	parser = ArgumentParser(description='VCF to anthracis genotypes')
	parser.add_argument('--mode', required=True,
		help='Mode to run in based on input files (vcf, bam, or vcf_parsnp)')
	parser.add_argument(
		'--vcf', nargs='+', type=str, required=False,
		help='VCF file(s) to genotype (Mapping MUST have been done using Ames Ancestor as a reference sequence)')
	parser.add_argument('--bam', nargs='+', type=str, required=False,
		help='BAM file(s) to genotype (Mapping MUST have been done using Ames Ancestor as a reference sequence)')
	parser.add_argument('--ref_id', type=str, required=False,
						help='Name of the reference in the VCF file (#CHROM column) or fasta file. Note that the Ames Ancestor has genotype 5.2 If all your strains return this genotype, it is likely you have specified the name of the refrence sequence incorrectly; please check your VCFs.')
	parser.add_argument('--phred', type=int, required=False, default=20,
						help='Minimum phred quality to count a variant call vs Ames as a true SNP (default 20)')
	parser.add_argument('--min_prop', type=float, required=False, default=0.1,
						help='Minimum proportion of reads required to call a SNP (default 0.1)')
	parser.add_argument('--ref', type=str, required=False,
						help='Reference sequence in fasta format. Required if bam files provided.')
	parser.add_argument('--output', type=str, required=False, default='genotypes.txt',
						help='Location and name for output file.')
	parser.add_argument('--samtools_location', type=str, required=False, default='',
						help='Location of folder containing samtools installation if not standard/in path.')
	parser.add_argument('--bcftools_location', type=str, required=False, default='',
						help='Location of folder containing bcftools installation if not standard/in path.')
	return parser.parse_args()

### Genotype SNP definitions

loci = [1771938, 1201909, 2212885, 2132895, 2019941, 3086718, 3366432, 3359543, 810017, 837700, 1271875, 857338, 4499321, 569295, 565120, 2332200, 2387683, 1776654]
snp_alleles = ['A', 'T', 'A', 'G', 'A', 'C', 'C', 'T', 'C', 'T', 'G', 'T', 'A', 'T', 'T', 'T', 'T', 'A']
groups = ['1.1', '1.2', '2.1', '2.2', '2.3', '3.1', '3.2', '3.3', '4.1', '4.2', '4.3', '5.1', '5.2', '5.3', '5.4', '6.1', '6.2', '6.3']

### QRDR SNP definitions (Dont need for B.anthracis)

#qrdr_loci = [2333762, 2333762, 2333750, 2333750, 2333751, 2333751, 3196469, 3196470, 3196458, 3196459]
#qrdr_snp_alleles = ['A', 'T', 'A', 'C', 'A', 'T', 'T', 'A', 'C', 'T']
#qrdr_groups = [' gyrA-S83F', ' gyrA-S83Y', ' gyrA-D87V', ' gyrA-D87G', ' gyrA-D87Y', ' gyrA-D87N', ' parC-S80R', ' parC-S80I', ' parC-E84G', ' parC-E84K']


# check if this SNP defines a QRDR group (Dont need for B.anthracis)
#def checkQRDRSNP(vcf_line_split, this_qrdr_groups, qrdr_proportions, args):
#	qrdr_snp = int(vcf_line_split[1])
#	if qrdr_snp in qrdr_loci:
#		i = qrdr_loci.index(qrdr_snp)
#		if float(vcf_line_split[5]) > args.phred:
#			print vcf_line_split
#			m = re.search("DP4=(\d+),(\d+),(\d+),(\d+)", vcf_line_split[7])
#			if m != None:
#				alt_read_count = int(m.group(3)) + int(m.group(4))
#				total_read_count = alt_read_count + int(m.group(1)) + int(m.group(2))
#				if float(total_read_count) == 0:
#					qrdr_snp_proportion = float(-1)
#				else:
#					qrdr_snp_proportion = float(alt_read_count) / total_read_count
#			else:
#				if vcf_line_split[4] != '.': # if the ALT is not '.' i.e. if the alt is not same as ref
#					try:
#						ad = vcf_line_split[9].split(':')[1].split(',') # get the AD ratio
#						alt_read_count = int(ad[1])
#						total_read_count = int(ad[0]) + alt_read_count
#						qrdr_snp_proportion = float(alt_read_count) / total_read_count
#					except IndexError:
#						qrdr_snp_proportion = float(-1)
#
#				else:
#					qrdr_snp_proportion = float(-1)	# set unknowns to negative so that we know this is not a real proportion
#
#			qrdr_snp_allele = vcf_line_split[4]
#			for position in xrange(0,10):
#				if (qrdr_snp == qrdr_loci[position]) and (qrdr_snp_allele == qrdr_snp_alleles[position]) and (qrdr_snp_proportion > args.min_prop):
#					this_qrdr_groups.append(qrdr_groups[position])  # Add QRDR SNP
#
#	return (this_qrdr_groups)


# check if this SNP defines a group
def checkSNP(vcf_line_split, this_groups, proportions, args):
	snp = int(vcf_line_split[1])
	if snp in loci:
		i = loci.index(snp)

		if float(vcf_line_split[5]) > args.phred:
			print (vcf_line_split)
			m = re.search("DP4=(\d+),(\d+),(\d+),(\d+)", vcf_line_split[7])
			if m != None:
				alt_read_count = int(m.group(3)) + int(m.group(4))
				total_read_count = alt_read_count + int(m.group(1)) + int(m.group(2))
				if float(total_read_count) == 0:
					snp_proportion = float(-1)
				else:
					snp_proportion = float(alt_read_count) / total_read_count
			else:
				if vcf_line_split[4] != '.': # if the ALT is not '.' i.e. if the alt is not same as ref
					try:
						ad = vcf_line_split[9].split(':')[1].split(',') # get the AD ratio
						alt_read_count = int(ad[1])
						total_read_count = int(ad[0]) + alt_read_count
						snp_proportion = float(alt_read_count) / total_read_count
					except IndexError:
						snp_proportion = float(-1)

				else:
					snp_proportion = float(-1)	# set unknowns to negative so that we know this is not a real proportion
			if snp_proportion > args.min_prop:
				this_allele = vcf_line_split[4]
				if this_allele == snp_alleles[i]:
					this_groups.append(groups[i])  # retrieve the group that this SNP defines
					proportions[groups[i]] = snp_proportion
	return (this_groups, proportions)

def checkSNPmulti(vcf_line_split, this_groups, args):
	snp = int(vcf_line_split[1])
	if snp in loci:
		i = loci.index(snp)
		strain = 0
		for gt in vcf_line_split[10:]:
			if (int(gt) == 1) and (vcf_line_split[4] == snp_alleles[i]):
				if strain in this_groups:
					this_groups[strain].append(groups[i])  # retrieve the group that this SNP defines
				else:
					this_groups[strain] = [groups[i]]
			strain += 1
	return this_groups


# sort groups into the three levels (primary, clade, subclade)
def parseGeno(this_groups, proportions):
	subclades = []
	clades = []
	primary = []
	for group in this_groups:
		level = len(group.split("."))
		if level == 5:
				subclades.append(group)
		if level == 4:
			subclades.append(group)
		if level == 3:
			subclades.append(group)
		elif level == 2:
			clades.append(group)
		elif level == 1:
			primary.append(group)

	# fix 1.2 nesting
	if ('1.2' in clades) and ('1.1' in clades):
		clades.remove('1.1')

	# fix 2.3, 2.2 nesting
	if ('2.1' in clades) and ('2.2' in clades):
		clades.remove('2.1')
	if ('2.2' in clades) and ('2.3' in clades):
		clades.remove('2.2')

	# fix 3.2, 3.3 nesting
	if ('3.1' in clades) and ('3.2' in clades):
		clades.remove('3.1')
	if ('3.1' in clades) and ('3.3' in clades):
		clades.remove('3.1')

	# fix 4.2, 4.3 nesting
	if ('4.1' in clades) and ('4.2' in clades):
		clades.remove('4.1')
	if ('4.2' in clades) and ('4.3' in clades):
		clades.remove('4.2')

	# fix 5.2, 5.3, 5.4 nesting
	if ('5.1' in clades) and ('5.3' in clades):
		clades.remove('5.1')
	if ('5.3' in clades) and ('5.4' in clades):
		clades.remove('5.3')

	# fix 6.2, 6.3 nesting
	if ('6.1' in clades) and ('6.2' in clades):
		clades.remove('6.1')
	if ('6.1' in clades) and ('6.3' in clades):
		clades.remove('6.1')

	 # fix clade relative to 5.2:
	if '5.2' in clades:
		clades.remove('5.2')  # anything NOT in 5.2 will have this SNP
	else:
		if len(clades) == 0:
			clades.append('5.2')  # anything with no clade, and 5.2 SNP not called, belongs in 5.2 with reference

	# store final genotype, to the lowest level available
	final_geno=''
	if primary:
		final_geno = ','.join(primary)
	if len(clades) > 0:
		final_geno = ','.join(clades)
	if len(subclades) > 0:
		final_geno = ','.join(subclades)

	# add proportion of reads supporting each of these groups
	p_prod = 1

	p_sub = []
	for group in subclades:
		if group in proportions:
			p_sub.append(str(round(proportions[group], 2)))
			p_prod = p_prod * proportions[group]

	p_cl = []
	for group in clades:
		if group in proportions:
			p_cl.append(str(round(proportions[group], 2)))
			p_prod = p_prod * proportions[group]

	p_pr = []
	for group in primary:
		if group in proportions:
			p_pr.append(str(round(proportions[group], 2)))
			p_prod = p_prod * proportions[group]

	# final call
	if final_geno=='':
		final_geno='None'

	info = final_geno + '\t'

	if 'A' in proportions:
		info += 'A'	 # annotate as 'A' to indicate this comes from assembled data and not reads
	else:
		info += str(round(p_prod, 2))  # indicate proportion of reads supporting this call

	# level calls
	info += '\t' + ','.join(subclades) + '\t' + ','.join(clades) + '\t' + ','.join(primary)

	# level proportions
	info += '\t' + ','.join(p_sub) + '\t' + ','.join(p_cl) + '\t' + ','.join(p_pr)

	# print(info)
	return info


# exception to raise if the command we try to run fails for some reason
class CommandError(Exception):
	pass


def run_command(command, **kwargs):
	'Execute a shell command and check the exit status and any O/S exceptions'
	command_str = ' '.join(command)
	logging.info('Running: {}'.format(command_str))
	try:
		exit_status = call(command, **kwargs)
	except OSError as e:
		message = "Command '{}' failed due to O/S error: {}".format(command_str, str(e))
		raise CommandError({"message": message})
	if exit_status != 0:
		message = "Command '{}' failed with non-zero exit status: {}".format(command_str, exit_status)
		raise CommandError({"message": message})


# main function
def main():
	args = parse_args()

	if (((args.mode == 'vcf') and args.vcf and args.ref_id) or ((args.mode == 'bam') and args.bam and args.ref and args.ref_id) or ((args.mode == 'vcf_parsnp') and args.vcf)):

		# Initialise output file and timestamp
		timestamp = datetime.datetime.fromtimestamp(time.time()).strftime('%Y.%m.%d_%H:%M:%S_')
		output_file = open(timestamp + args.output , 'w')

		# GENERATE VCFS (1 per strain) FROM BAMS

		if args.mode == 'bam':

			with open(args.ref, 'r') as fasta_file: # create SAMtools compatible fasta file
				sequences = fasta_file.read()
				for sequence in sequences.split('>'):
					if args.ref_id in sequence:
						new_header = '>' + args.ref_id
						replacement_index = sequence.find('\n')
						sequence = new_header + sequence[replacement_index:]
						with open('temp_reference.fasta', 'w') as temp_fasta_file:
							temp_fasta_file.write(sequence)


			vcfFiles = []

			# coordinates in zero-base, half-open for SAMtools compatible bed file
			ordered_loci = list(loci)
			sorted(ordered_loci)
			temp_bed_file = open(args.ref_id + '.bed', 'w')	 # create temporary bed file for SAMtools
			for locus in ordered_loci:
				temp_bed_file.write(
					args.ref_id + '\t' + str(locus - 1) + '\t' + str(locus) + '\n')	 # write bed file from matrix
			temp_bed_file.close()  # close bedFile

			run_command(['samtools', 'faidx', 'temp_reference.fasta'])	# index fasta file

			for bam in args.bam:
				print ('bam files supplied, generating vcf file for ' + bam)
				if os.path.exists(bam + '.bai') == False: # index bam file if indexed bam not provided
					run_command([args.samtools_location + 'samtools', 'index', bam])

				run_command(
					[args.samtools_location + 'samtools', 'mpileup', '-q', str(args.phred), '-ugB', '-f', 'temp_reference.fasta',
				 	'-l', args.ref_id + '.bed', bam, '-o', bam[:-4] + '.output', '-I']) # detect SNPs

				run_command(
					[args.bcftools_location + 'bcftools', 'call', '-c', bam[:-4] + '.output', '-o', bam[:-4] + '.vcf']) # generate vcf files
				run_command(['rm', bam[:-4] + '.output'])

				vcfFiles.append(bam[:-4] + '.vcf')	# supply generated vcf file to script

			run_command(['rm', args.ref_id + '.bed']) # remove temp files
			run_command(['rm','temp_reference.fasta'])
			run_command(['rm', 'temp_reference.fasta.fai'])

			args.vcf = vcfFiles


		# PRINT OUTPUT HEADER

		if args.mode == 'bam':
			output_file.write('\t'.join(
				['File', 'Classification\n']))
		elif args.mode == 'vcf':
			output_file.write('\t'.join(
				['File', 'Classification\n']))
		else:
			output_file.write('\t'.join(
				['File', 'Classification\n']))


		# PARSE MAPPING BASED VCFS (1 per strain)

		if (args.vcf and (args.mode != 'vcf_parsnp')):
			print('vcf mode')
			for vcf in args.vcf:
				snp_count = 0
				this_groups = []  # list of groups identified by defining SNPs
				#this_qrdr_groups = [] # list of QRDR SNPs found
				proportions = {}  # proportion of reads supporting each defining SNP; key = group, value = proportion
				#qrdr_proportions = {}  # proportion of reads supporting each defining SNP; key = group, value = proportion

				# read file
				(file_name, ext) = os.path.splitext(vcf)

				if ext == '.gz':
					f = gzip.open(vcf, 'r')
				else:
					f = open(vcf, 'r')

				any_ref_line = 0

				for line in f:
					if not line.startswith('#'):
						x = line.rstrip().split()
						if not line.strip() == '':
							snp_count = snp_count + 1
						if x[0] == args.ref_id:
							# parse this SNP line
							any_ref_line = 1
							(this_groups, proportions) = checkSNP(x, this_groups, proportions, args)

							#Remove QRDR groups (SB)
							#this_qrdr_groups = checkQRDRSNP(x, this_qrdr_groups, qrdr_proportions, args)

				f.close()

				# remove qrdr groups (SB)
				# print "qrdr groups".join(qrdr_groups)
				if any_ref_line > 0:
					info = parseGeno(this_groups, proportions)

					if args.bam:
						# output_file.write(vcf + '\t' + info + '\t' + str(snp_count) + '\t' + ','.join(this_qrdr_groups) + '\n')
						output_file.write(vcf + '\t' + info.split('\t')[0] + '\n')# + '\t' + str(snp_count) + '\t\n')
					else:
						# output_file.write(vcf + '\t' + info  + '\t' + ','.join(this_qrdr_groups) + '\n')
						output_file.write(vcf + '\t' + info.split('\t')[0] + '\n')#  + '\t' + '\n')
				else:
					output_file.write(
						vcf + '\tNo SNPs encountered against expected reference. Wrong reference or no SNP calls?\n')

		# PARSE PARSNP VCF (multiple strains)

		if args.mode == 'vcf_parsnp':

			if not args.ref_id:
				args.ref_id = '1'

			for vcfm in args.vcf:

				# read file
				(file_name, ext) = os.path.splitext(vcfm)

				if ext == '.gz':
					f = gzip.open(vcfm, 'r')
				else:
					f = open(vcfm, 'r')

				any_ref_line = 0

				this_groups = {}  # list of groups identified by defining SNPs, key = strain id (column number)
				strains = []

				for line in f:
					x = line.rstrip().split()
					if x[0] == '#CHROM':
						strains = x[10:]
					if not line.startswith('#'):
						if x[0] == args.ref_id:
							any_ref_line = 1  # parse this SNP line
							this_groups = checkSNPmulti(x, this_groups, args)


				f.close()

				# collate by strain
				if any_ref_line > 0:
					# print(any_ref_line)
					for strain in this_groups:
						info = parseGeno(this_groups[strain], ['A'])
						output_file.write(strains[strain] + '\t' + info + '\n')
				else:
					output_file.write(strains[
						strain] + '\tNo SNPs encountered against expected reference. Wrong reference or no SNP calls?\n')

		output_file.close()
	else:
		print ('Missing or incomplete input parameters, please check these and try again.')

# call main function
if __name__ == '__main__':
	main()
